# Infectome Analysis of Small Mammals in Southern China Reveals Ecological Associations and Emerging Threats from Diverse Pathogens

**DOI:** 10.1101/2025.02.09.637073

**Authors:** Genyang Xin, Daxi Wang, Xin Zhang, Qingquan Cen, Minwu Peng, Yuqi Liao, Jing Wang, Shijia Le, Jinxia Cheng, Wei-chen Wu, Xin Hou, Gengyan Luo, Qinyu Gou, Jianbin Kong, Zhu Pan, Dongqing Li, Shipei Gan, An Chen, Hailong Zhao, Peibo Shi, Zirui Ren, Weiqi Zhao, Jiajun Liu, Penghui Jia, Changyun Sun, Wenqing Lin, Jun Wu, Guopeng Kuang, Jingdiao Chen, Junhua Li, Edward C. Holmes, Ziqing Deng, Mang Shi

**Affiliations:** National Key Laboratory of Intelligent Tracking and Forecasting for Infectious Diseases, School of Medicine, Shenzhen Campus of Sun Yat-sen University, Sun Yat-sen University, Shenzhen, China; State Key Laboratory for Biocontrol, School of Medicine, Shenzhen Campus of Sun Yat-sen University, Sun Yat-sen University, Shenzhen, China; Shenzhen Key Laboratory for Systems Medicine in Inflammatory Diseases, Shenzhen Campus of Sun Yat-sen University, Sun Yat-sen University, Shenzhen, China; BGl Research, Beijing, China; Shenzhen Key Laboratory of Unknown Pathogen Identification, BGl Research, Shenzhen, China; Institute of Pathogenic Microbiology, Center for Disease Control and Prevention of Guangdong Province, Guangzhou, China; Guangdong Institute of Plague Control and Prevention, Zhanjiang, China; Lianjiang Center for Disease Control and Prevention, Lianjiang, China; College of Life Sciences, University of Chinese Academy of Sciences, Beijing, China; Institute of Communicable Disease Control and Prevention, Center for Disease Control and Prevention of Guangdong Province, Guangzhou, China; Institute of Disinfection and Vector Control and Prevention, Center for Disease Control and Prevention of Guangdong Province, Guangzhou, China; Yunnan Provincial Key Laboratory for Zoonosis Control and Prevention, Yunnan Institute of Endemic Disease Control and Prevention, Kunming, China; Institute of Parasitic Disease Control and Prevention, Center for Disease Control and Prevention of Guangdong Province, Guangzhou, China; School of Medical Sciences, The University of Sydney, Sydney, Australia.; Laboratory of Data Discovery for Health Limited, Hong Kong SAR, China

## Abstract

Small mammals harbor a diverse array of zoonotic pathogens. To date, however, metagenomic surveys of these species have primarily focused on viral diversity, with limited attention paid to bacteria and eukaryotic pathogens. Additionally, the ecological determinants of pathogen diversity within these mammals have not been systematically examined. Herein, we employed a metatranscriptomics approach to survey the pathogen infectome of 2,408 individual samples, representing lung, liver, and gut tissues from 858 animals collected throughout Guangdong province, China, with a study design that accounted for host species, tissue, season, and geographic location. We identified 76 pathogen species, including 29 RNA viruses, 12 DNA viruses, 5 bacteria, and 30 eukaryotic pathogens, 33 of which were newly discovered. Tissue distribution analysis revealed distinct organotropisms, suggesting varied transmission routes, while host distribution analysis showed that each animal carried an average of one pathogen, with 10 pathogens widely distributed among mammalian orders. Our characterization of the geographic and seasonal patterns revealed that pathogen richness was primarily influenced by region, host species, and season, while pathogen composition was largely shaped by host genetic distance. Collectively, these data provide the first comprehensive insight into the dynamics of the pathogen infectome in these key mammalian disease reservoirs, highlighting the major factors driving pathogen diversity and transmission.

## Introduction

Small mammals, including rodents (order Rodentia) and shrews (order Eulipotyphla), are major reservoirs of zoonotic pathogens that pose significant risks to human health. These pathogens include hemorrhagic fever viruses (such as arenaviruses and hantaviruses), that can have a mortality rate of up to 81% ^1^; plague and fever-causing bacteria (e.g., *Yersinia pestis*, *Rickettsia*, and *Leptospira*), which can lead to serious disease but are sporadically detected in China ^2–9^; and various endoparasites (e.g., *Toxoplasma*), that can infect humans but depend on rodents for part of their life cycle ^10^. Beyond the pathogens they carry, the public health risk posed by small mammals is amplified by their close contact with humans ^11–14^. Indeed, despite significant improvements in hygiene and sanitation over the past two decades, pathogens associated with small mammals can be transmitted through direct animal contact, via their excretions, contaminated water and food, or through blood-sucking fleas and ticks ^3,9,15–18^.

As small mammals can serve as reservoirs for important pathogens and sometimes closely interact with humans, modern pathogen discovery has increasingly focused on these animals. The goal of these studies is to identify a broader diversity of potential pathogens, although most have concentrated on viruses. Earlier studies using PCR assays with primers designed from conserved viral genome regions led to the discovery of numerous new rodent- and shrew-associated viruses within all major families of mammalian viruses, often forming host-associated virus clusters and filling gaps in diversity ^19–27^. The known diversity of viruses in these groups has further expanded with the use of viral-particle enrichment metagenomics ^28–30^ and meta-transcriptomics ^31–34^. For example, a total of 206 virus species were identified in a study involving 3055 individuals from 50 rodent species, establishing the core virome of these mammals in China ^28^. Similarly, a meta-transcriptomic survey of shrew lung virome revealed more novel than known viruses^35^. Notably, some studies also uncovered pathogens related to those that cause disease in humans, such as Mojiang virus – a close relative of the Hendra and Nipah viruses discovered in an abandoned mine shaft ^28,36^. Collectively, these studies provide a broad-scale investigation of the virome in rodents and shrews, expanding our understanding of viral diversity and identifying potential emerging human pathogens.

Large-scale sampling and sequencing has also greatly advanced studies of pathogen ecology and evolution ^34,37–39^. These studies have compared viral diversity across regions and host species, revealing strong host-virus associations, with geographic factors playing a secondary role in shaping virus diversity ^28,35,40^. Importantly, instances of cross-species virus transmission have also been documented, occurring between different host species and, less frequently, across host families and orders^41,42^. Despite these findings, ecological comparisons have generally been conducted at a broad scale, frequently lacking study designs that account for host species, geographic location, temporal distribution, and adequate replication across variables. Moreover, many studies have relied on pooled samples, which sometimes lead to the mixing of host species within a single sample, greatly complicating the analysis of cross-species virus transmission. Additionally, the focus on viral diversity has overshadowed investigations into the presence and abundance of other pathogen types, such as bacteria and eukaryotic pathogens, leaving significant gaps in our understanding of their ecological roles and interactions.

Herein, we describe a large-scale survey of small mammals and their associated pathogens in Guangdong province in southern China. Guangdong is located in the subtropical climate zone and boasts high biodiversity. Historically, Guangdong has also been an epicenter for the emergence SARS and other infectious disease outbreaks of importance ^43^. Additionally, as a major gateway for both the domestic and international trade, Guangdong plays a significant role in infectious disease dissemination and spread. Our study comprised a broad geographic survey conducted in winter, as well as a year-long monthly survey in two cities, encompassing both domestic and rural small mammal populations. Meta-transcriptomic sequencing was performed on individual lung, spleen, and gut samples from single animals. With these data in hand we systematically compared the full spectrum of pathogens – RNA viruses, DNA viruses, bacteria, and eukaryotic pathogens – across different organs, host species, geographic locations, and seasons. Our findings provided valuable insights into pathogen diversity under varying conditions and revealed key factors driving pathogen evolution and transmission.

## Results

### Overview of host and pathogen diversity

We conducted a systematic survey of small mammals and their associated pathogens (i.e., those microbes, including viruses, known or likely to be associated with disease) across Guangdong province, China (Fig. 1a). The survey comprised two approaches: one covering nine regions across Guangdong sampled during the winter season, and the other spanning different months throughout the year, focusing on two locations – Anpu and Zhanjiang in southwestern Guangdong province. Sampling sites in each region included both agricultural and residential areas. In total, 858 individual animals were collected. COX1 analysis identified nine species across six genera: *Rattus*, *Bandicota*, *Berylmys*, *Mus*, *Niviventer*, and *Suncus*, belonging to two mammalian orders, Rodentia and Eulipotyphla (Fig. 1b). *Rattus norvegicus* was the most commonly sampled species (*N* = 240), followed by *Bandicota indica* (*N* = 195), *Rattus tanezumi* (*N* = 151), *Suncus murinus* (*N* = 118), and *Rattus losea* (*N* = 92). While species composition varied significantly across geographic locations, it remained relatively stable across different seasons (Fig. 1c and d, Table S1).

**Figure 1.**
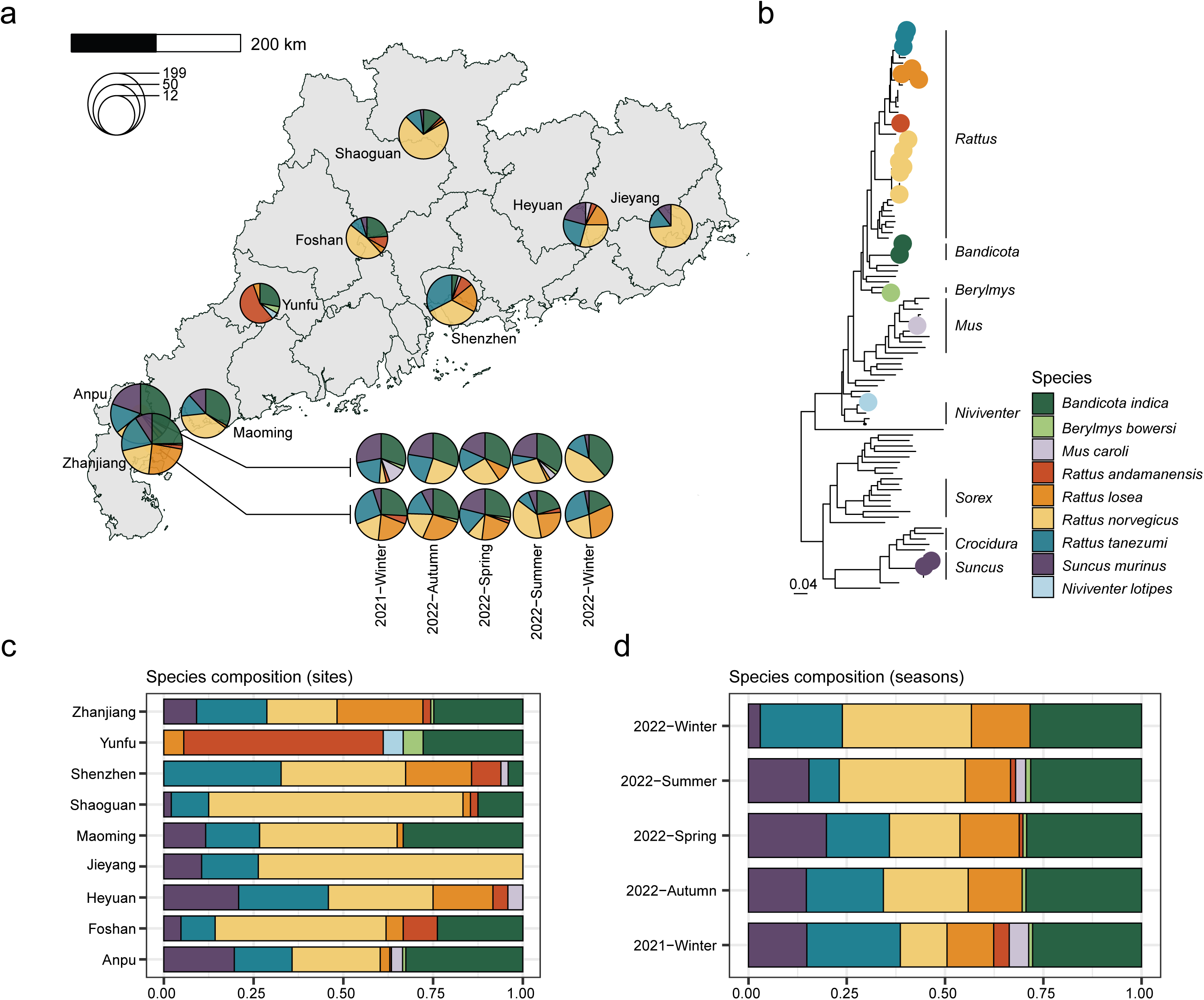
Sample distribution of small mammals in this study. (a) The pie charts show the distribution of host species across the sampling locations. The size of each pie chart reflects the sample size, and the colors represent the species composition. (b) Phylogenetic relationships of the sampled species (denoted by solid circles) and their related mammalian species. (c) Distribution of host species by sampling site. (d) Seasonal distribution of host species in Anpu and Zhanjiang, Guangdong province.

For all 858 animals sampled, we performed independent meta-transcriptomic sequencing on three tissues—lung, liver, and gut—resulting in the production of 2,408 sequencing libraries and a total of 10.2 Tbp of data. The median sequencing depth per library was 42.85 Gbp. From this data set, we identified 80 species of mammalian RNA viruses, 20 mammalian DNA viruses, 5 bacterial pathogens, and 30 eukaryotic pathogens, spanning 15 viral, 4 bacterial, and 17 eukaryotic families. Due to the unknown hosts of members from the families *Picobirnaviridae* and *Anelloviridae*, they were excluded as potential pathogens. Consequently, the pathogen infectome of the sampled mammalian species, based on the analyzed tissues, comprised 76 microbial species, namely, 29 RNA viruses, 12 DNA viruses, 5 bacterial, and 30 eukaryotic pathogens. (Fig. 2, S1, S2, S3, Table S2 & S3)

**Figure 2.**
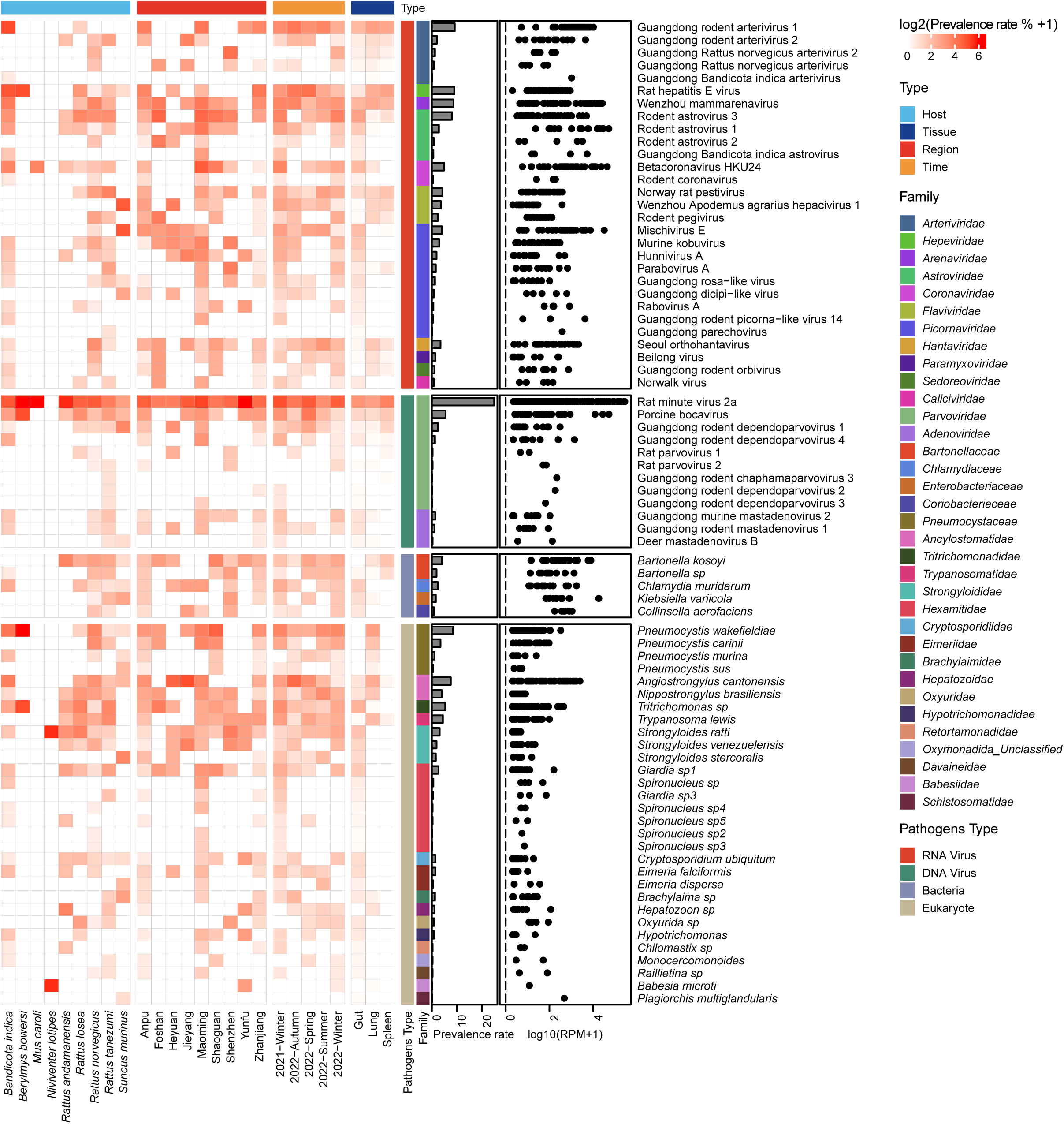
Diversity and prevalence of mammal-associated pathogens in sampled small mammals. The heatmap (left panel) illustrates the distribution of pathogens across host species, sampling sites, seasons, and tissues, with color intensity indicating the log-transformed prevalence rate of each pathogen. The histogram (middle panel) shows the overall prevalence rate of pathogens, while the scatter plot (right panel) displays the abundance levels (log2(prevalence rate %+1)) of pathogens.

**Figure 3.**
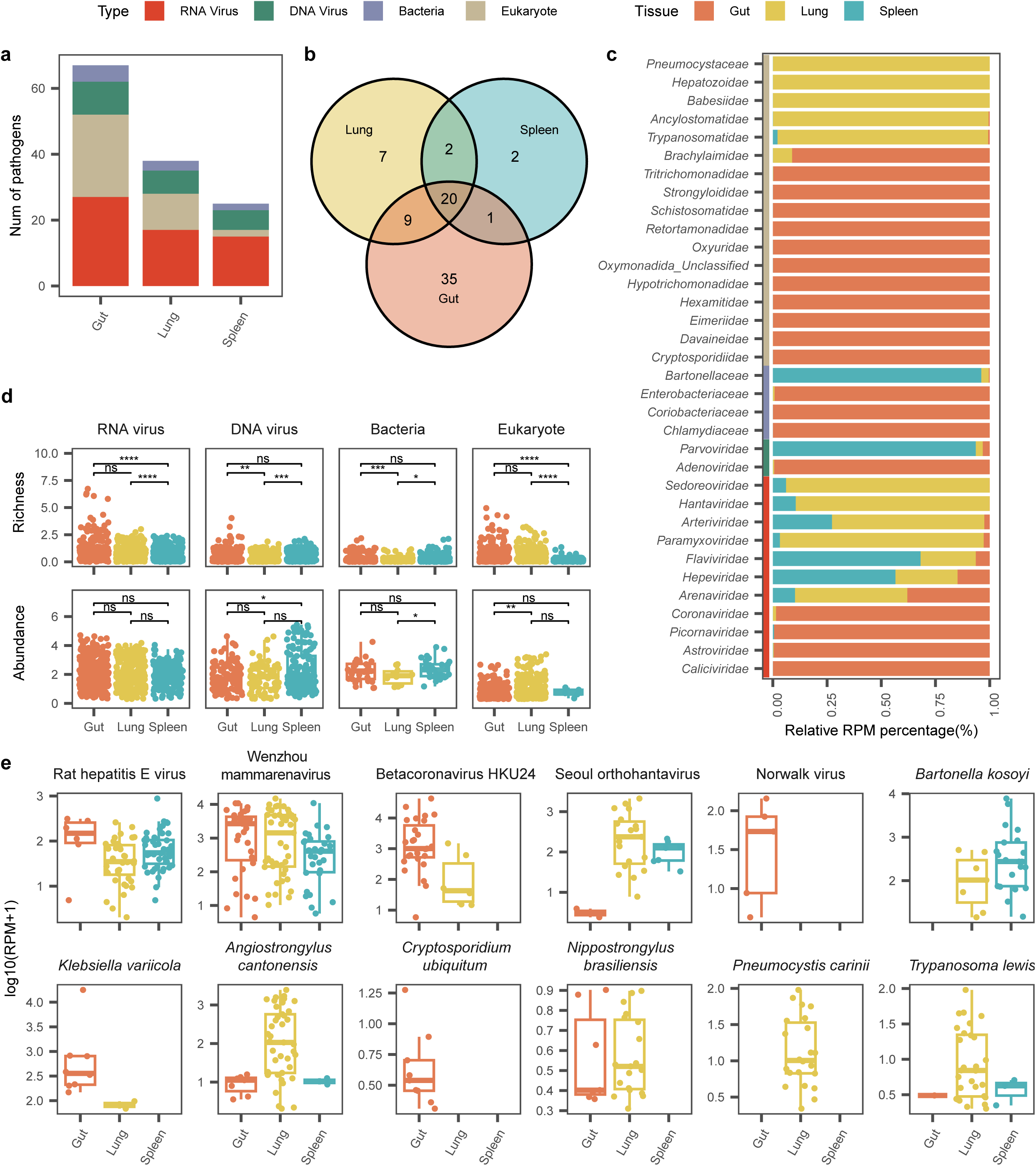
Organotropism of mammal-associated pathogens. (a) Bar graph showing number of RNA viruses, DNA viruses, bacteria, and eukaryotes detected in the gut, spleen, and lung. (b) Venn diagram showing the overlap of pathogen species between different tissues. (c) Comparisons of pathogen abundance across three tissues. (d) Comparisons of pathogen richness (top panel) and abundance (bottom panel) across three tissues. (e) Organotropism of the 12 most abundant zoonotic pathogens.

### The pathogen infectome of small mammals in Guangdong province

Among the 100 mammalian viruses discovered, 69 were newly identified species, with the remainder representing existing species (Fig. S4, S5). More novel viral species were identified from the RNA virus families *Arteriviridae* (N = 5) and DNA virus family *Parvoviridae* (N = 6). Notably, a novel orbivirus, provisionally named Guangdong rodent orbivirus, formed a distinct sister lineage, displaying significant divergence from known orbiviruses. However, that it was identified in lung and spleen at a relatively high abundance (up to 723 RPM) suggests it is likely to be a mammalian virus. Additionally, novel members of the bacterial genus *Bartonella* and eukaryotic genera such as *Giardia*, *Spironucleus*, *Brachylaima*, and *Hepatozoon* were identified and confirmed through analyses of marker genes, occupying distinct positions on the phylogenetic trees (Fig. 2, Table S3).

At the level of host species, the highest prevalence among RNA viruses was observed for Guangdong rodent arterivirus 1 (9.15%). Zoonotic pathogens such as Rat hepatitis E virus (8.99%), Wenzhou mammarenavirus (8.68%), Betacoronavirus HKU24 (4.89%), and Seoul orthohantavirus (3.47%) were also found at relatively high prevalence. Among DNA viruses, parvoviruses showed a relatively high prevalence, with Rat minute virus 2a exceeding 20%, while adenoviruses were less frequently detected (2.83%) (Fig. 2, Table S3).

Notably, our study revealed a high prevalence (40.82%) of eukaryotic pathogens. Of these, *Pneumocystis* (9.96%), a parasitic fungus commonly found in mammalian lungs, and *Angiostrongylus cantonensis* (Rat lungworm, 7.57%), a nematode parasite of the lung, were among the most prevalent. Other gut nematodes such as *Nippostrongylus brasiliensis* (3.94%) and Strongyloides (5.92%) were also abundant. Protozoan parasites such as *Tritrichomonas* (4.9%), *Trypanosoma* (4.04%), and *Giardia* (2.87%) were also relatively commonplace. Additionally, zoonotic human pathogens such as *Cryptosporidium ubiquitum*, a fungal species, was identified in nine hosts (1.42%), and *Babesia microti*, a blood-borne parasitic species, was found in the lungs of a *Niviventer lotipes*. In contrast, we detected fewer bacterial pathogens, although *Bartonella* (5.99%) and *Chlamydia* (2.37%) had relatively high prevalence rate (Fig. 2, Table S3).

### Tissue-specific distribution of pathogens

The distribution and abundance of mammal-associated pathogens were assessed across three tissues: gut, lung, and spleen. The gut harbored the highest pathogen diversity (67 species), followed by the lung (38 species) and spleen (25 species) (Fig. 3a). Overall, 25.6% (20/78) of the pathogens were shared among all three tissues, of which 13 were RNA viruses, including Wenzhou virus, Rat hepatitis E virus, Beilong virus and *Orthohantavirus seoulense* (Fig. 2, 3b and 3c). Other pathogens similarly displayed distinct organotropisms. For example, viruses from the *Picornaviridae*, *Coronaviridae*, and *Caliciviridae* were more prevalent in the gut, while DNA viruses from *Parvoviridae* and bacteria from *Bartonella* were enriched in the spleen. Interestingly, several eukaryotic parasites, including *Pneumocystis*, *Angiostrongylus*, *Trypanosoma* and *Babesia*, were primarily detected in the lungs (Fig. 3c). The gut generally exhibited the greatest pathogen diversity across all categories, with the exception of DNA viruses, for which the spleen harbored a similar level of diversity with gut but with greater abundance. However, when considering pathogen abundance, the gut did not demonstrate the same dominance as observed for pathogen diversity. Indeed, in the case of eukaryotic pathogens, the lungs harbored a higher abundance than the gut. Finally, analysis of the 12 most abundant zoonotic pathogens across the tissues revealed that 10 were likely transmitted via the respiratory route, nine through the fecal-oral route, and six via the circulatory system, with *Bartonella kosoyi* likely transmitted by arthropod vectors (Fig. 3e).

### Distribution and transmission of pathogens among different hosts

We first assessed pathogen richness in each host species examined. On average, individual animals carried a median of one pathogen species (range: 0-12) across the three tissue types sampled (Fig. 4a). Notably, 30.3% of the individuals were pathogen-free, based on the detection threshold set in this study. Comparisons across different host species revealed significant variability in pathogen richness, with *Bandicota indica* and *Rattus norvegicus* carrying significantly more pathogens per individual compared to *Suncus murinus* and *Rattus losea* (Fig. 4b).

**Figure 4.**
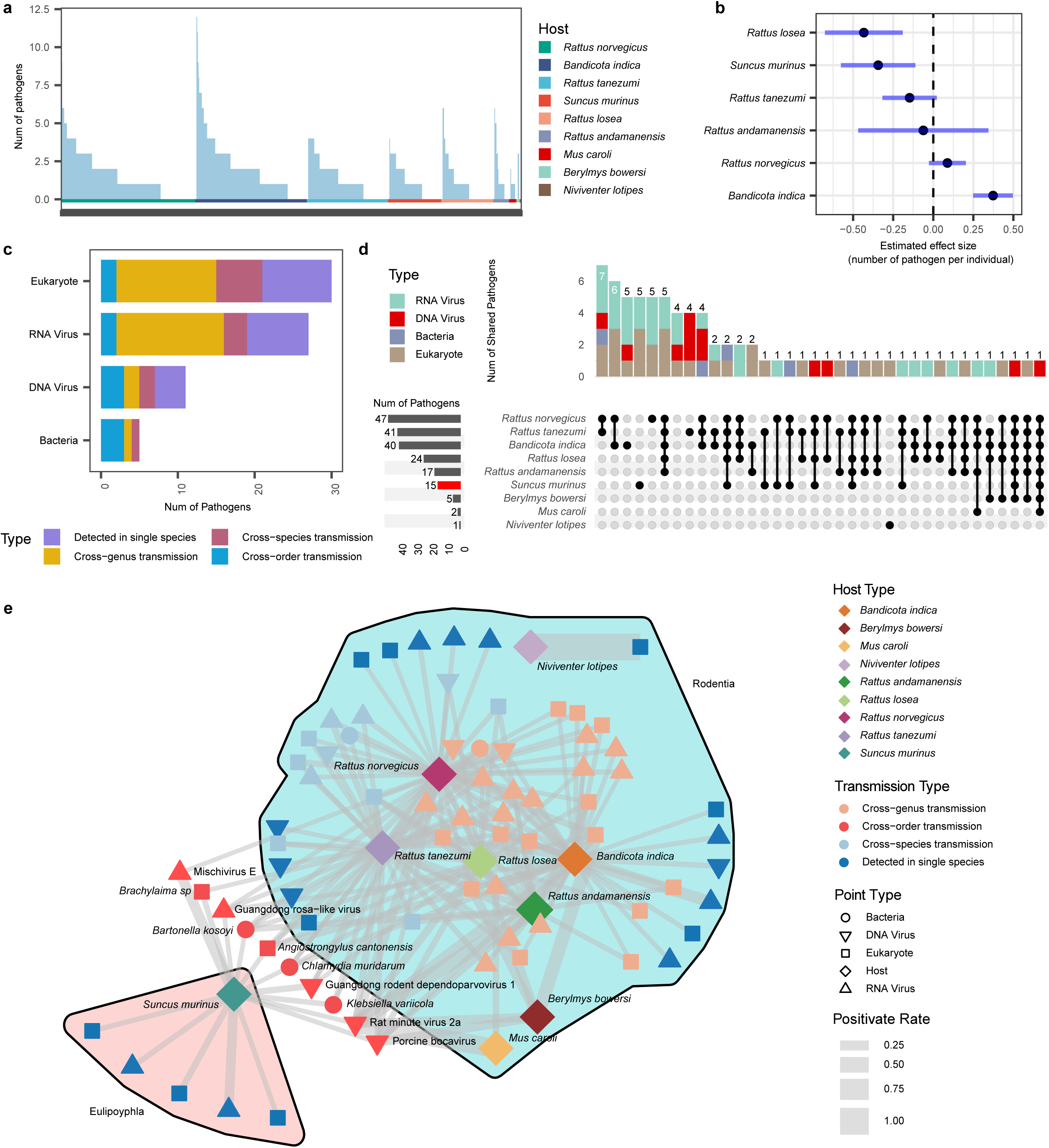
Pathogen diversity and transmission among different hosts. (a) Number of pathogens carried by each individual host. The color bar at the bottom indicates host species. (b) Estimated effect size of mammalian species on pathogen richness per individual, presented with estimated values and 95% confidence intervals (CI). (c) Number of pathogens shared across species, genera, families, and orders of different mammalian hosts. Four pathogen types are represented using distinct colors. (d) UpSet graph illustrating the number of pathogens shared between different host species. (e) Virus sharing network. Nodes represent hosts or virus species, colored by host species and cross-species transmission potential, and shaped according to pathogen types. Line thickness between nodes reflects the positivity rate in less prevalent hosts, with the network divided into two subnets based on host order. External nodes highlight pathogens with potential for cross-order transmission.

We further examined pathogen transmission across host species, genera, families, and orders. Cross-species transmission appeared to be the rule rather than the exception, with 65.8% of the pathogens detected showing the ability to cross host species barriers and 12.7% able to infect hosts from multiple mammalian orders (Fig. 4 c, d). Notably, all the bacterial species identified were capable of infecting multiple host species. Pathogens present in multiple host orders represent those with the broadest host ranges and hence may be of increased zoonotic potential. In total, 10 cross-order pathogens were discovered, including two RNA viruses, three DNA viruses, two bacterial species, and three eukaryotic pathogens (Fig. 4e). Importantly, eight of these cross-order pathogens were found in more than two species, with similarly high prevalence rates (Fig. S6), suggesting a heightened risk to public health.

### Epidemiological correlates of the total infectome

We analyzed the factors influencing pathogen diversity in terms of richness (Fig. 5a) and species composition (Fig. 5b). The largest variation in viral richness was explained by the geographic region where samples were collected (11.4%, Fig. 5c), followed by host species identity (6.8%), sampling month (6.5%, Fig. 5d), and environmental factors (3.2%, Fig. 5e). Specifically, spring and winter seasons, along with the towns of Jieyang and Maoming, were associated with higher pathogen richness (Fig. 5c and d). However, 72.4% of the total variation remained unexplained (Fig. 5a). Pathogen composition, assessed by shared microbial species between sample groups, was most strongly influenced by host phylogenetic distance, explaining 11.1% of the variation (Fig. 5b). Other factors showed no significant effect. As expected, there was a significant negative correlation between host genetic distance and shared pathogen species (Fig. 5f).

**Figure 5.**
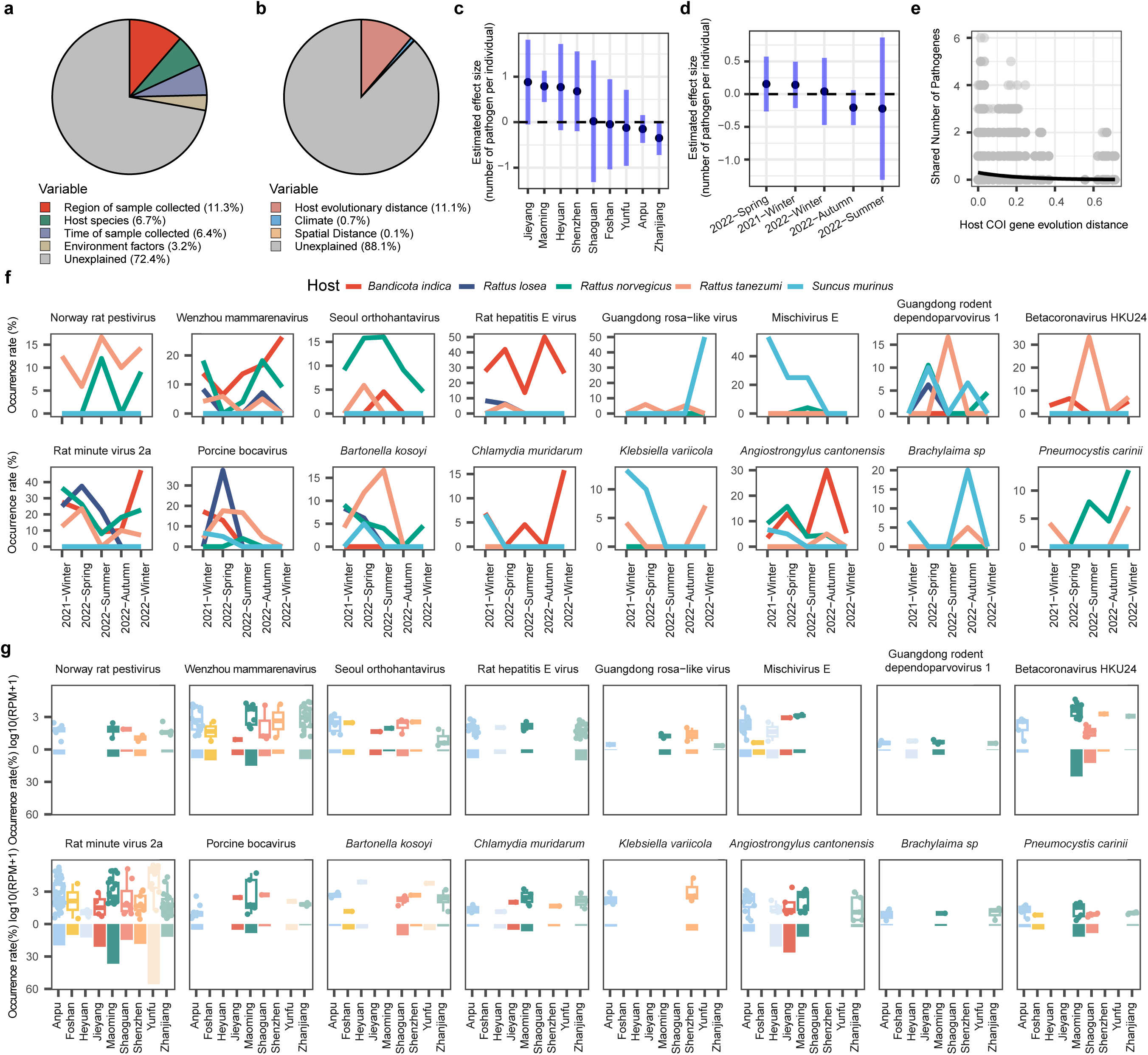
Ecological associations of the pathogens circulating in small mammals. (a) Relative contribution of sampling location, host species, sampling season, and meteorological data to pathogen richness in each individual animal, quantified by the explained deviance in the best model structures (ΔAIC <2) using generalized linear models (GLMs). (b) Relative contribution of host evolution distance, climate, and spatial distance to share number of pathogens between individuals. (c) Estimated effect size of sampling location on pathogen richness per individual, presented with estimated values and 95% confidence intervals (CI). (d) Estimated effect size of sampling season on pathogen richness per individual, presented with estimated values and 95% confidence intervals (CI). (e) Relationship between the number of shared pathogens and host phylogenetic distance (patristic distance based on mitochondrial COX1 gene phylogeny) among pairs of animal individuals. (f) Seasonal variation in the occurrence rate of 16 high zoonotic potential pathogens. (g) Geographic comparison of prevalence rates and abundance levels for 16 high zoonotic potential pathogens.

We then analyzed the seasonal and geographical dynamics of pathogens circulating in small mammals across Guangdong province. In terms of seasonal prevalence (Fig. 5g), several pathogens showed winter peaks, including Wenzhou mammarenavirus, *Chlamydia muridarum*, and *Pneumocystis carinii*, while others, like Rat hepatitis E virus and *Angiostrongylus cantonensis*, peaked in autumn. Notably, a number of pathogens, including Seoul orthohantavirus, Betacoronavirus HKU24, and *Bartonella kosoyi*, had higher prevalence during the summer. Geographically, zoonotic pathogens such as Wenzhou mammarenavirus, Seoul orthohantavirus, and *Bartonella kosoyi* were widely distributed, present in the majority of sampled regions. Wenzhou mammarenavirus, Betacoronavirus 1, and *Pneumocystis carinii* were most prevalent in Maoming, whereas parasites *Angiostrongylus cantonensis* was more frequently detected in Jieyang and Heyuan (Fig. 5h).

## Discussion

We conducted a large-scale surveillance of a broad spectrum of pathogens, including RNA viruses, DNA viruses, bacteria, fungi, and parasites. Previous infectome studies have largely taken a smaller scale perspective. For instance, studies have mapped the full spectrum of pathogens in the human respiratory system ^44,45^, highlighted shifts in opportunistic pathogens and commensal microbes in the respiratory tract following SARS-CoV-2 infection and their link to differing clinical outcomes^45^, and demonstrated the importance of pathogen panels over single-pathogen models in explaining diseases in pigs ^46^. Building on these insights, our study employed the same meta-transcriptomics approach with a study design encompassing diverse host ranges, geographic regions, seasonal variations, and tissue/transmission types, along with individual-level sequencing. In doing so we unveiled, for the first time, the macro-ecological patterns of diverse pathogens within small mammals that are key reservoirs for various infectious diseases ^38,47^. Specifically, we both identified and characterized them in detail, uncovering insights into their diversity, prevalence, seasonal trends, organotropism, and potential for cross-species transmission. This metagenomics-based ecological and epidemiological framework represents a powerful tool that can be readily adapted to study pathogens in other organisms or environments, offering valuable data for understanding and mitigating risks of pathogen transmission.

Our results revealed that the zoonotic risks associated with these small mammals are significant, although most of the pathogens identified were known species. For instance, Guangdong province has a relatively high incidence of Hemorrhagic Fever with Renal Syndrome, with 93-328 cases reported each year between 2015 and 2021 ^48^. Herein, we reveal that the causative virus, Seoul orthohantavirus, is widely distributed across the region and is among the most prevalent zoonotic pathogens in Guangdong, with a prevalence rate of 3.47%. In contrast, while Wenzhou mammarenavirus is also highly prevalent in rodent populations (8.67%), reported human cases remain rare. This virus was initially identified in various small mammals, including rodents and shrews, in Wenzhou, Zhejiang^49^. Since then, it has been widely detected across Southern China and Southeast Asia^28,31,50–54^. Despite its broad prevalence in animals, human infections appear uncommon, likely reflecting limited adaptation of the virus to humans. Alternatively, the virus may cause mild, nonspecific symptoms that go unrecognized and are not routinely screened. Supporting this, a study from Cambodia revealed that RT-PCR assays typically only detect the virus in patients with respiratory symptoms^52^. A serological survey of the same population found a 13.23% IgG seroprevalence among healthy individuals ^52^, suggesting undiagnosed or asymptomatic infections are more widespread.

One surprising finding was the notably high load of eukaryotic pathogens, all of which were confirmed through phylogenetic analyses based on marker genes. This suggests that these animals are highly susceptible to these parasites, which have the potential to infect a variety of mammalian hosts, including humans^55–57^. Consequently, these animals may act as key maintenance or amplifying hosts for these parasites^56^. This raises major public health concerns for two main reasons: (i) unlike viruses, eukaryotic pathogens have the potential to infect many groups of mammals including humans^58^, and (ii) the transmission routes to humans are diverse, including direct contact, aerosolized particles, contaminated food or water, and arthropod vectors^58,59^. Additionally, the particularly high diversity of eukaryotic pathogens underscores the importance of adopting new surveillance approaches for these pathogens. Previous surveillance programs often did not include these animal species, and since it is challenging to detect these pathogens directly from the environment or intermediate hosts, monitoring their prevalence in mammalian hosts offers a more effective means of surveillance.

Our study revealed that the richness of pathogens in rodents is influenced by various ecological and biological factors. We observed significant seasonal, geographic, and host-related variations in pathogen diversity. Notably, the highest pathogen richness was found in *Bandicota indica*. This is likely due to wild rodents having broader ecological interactions, occupying diverse habitats, and interacting with larger host communities that facilitate pathogen transmission. In contrast, house rats, which inhabit human environments, typically have a more restricted living range and lower species diversity, limiting opportunities for virus transmission^14,38,56,60^. We also observed seasonal variations in pathogen diversity, with higher overall diversity detected during winter and spring. This pattern may result from increased rodent population density following the autumn food harvest, coupled with a 2–14 weeks lag before pathogen diversity peaks ^38,56,61–63^. However, this trend was not highly significant, likely due to the relatively small sample size. Alternatively, the observed patterns may reflect the fact that different pathogens peak in different seasons, which can influence overall trends.

In addition to pathogen richness, we compared pathogen composition across different samples. This revealed that host species is the most important factor shaping pathogen composition, consistent with many other studies characterizing viruses circulating in small mammals ^28,31,40^. Despite this, our study identified several pathogens capable of crossing host species, and, to a lesser extent, host orders. Of particular interest are those pathogens that can jump between host orders. Specifically, we identified two RNA viruses, three DNA viruses, three bacteria, and two parasites that are capable of transmission across different host orders. Among these, *Angiostrongylus cantonensis*, *Klebsiella variicola*, and *Bartonella kosoyi* are well-known zoonotic pathogens with broad host ranges, while Mischivirus E, *Brachylaima sp*, and Porcine bocavirus are less recognized for their ability to infect both animals and humans or unclear pathogenic nature. However, that these viruses can infect both rodents and shrews suggests that they may be host generalists and hence are at threat of zoonotic spillovers.

Our study has several limitations. First, while it represents the largest individual-level sample size to date, it remains limited with respect to the total number of individuals involved. Certain species, such as *Mus caroli*, *Niviventer lotipes*, and *Berylmys bowersi*, are still under-represented, although the total numbers of these animals may be small. Second, our study may underestimate the pathogen diversity, particularly bacteria and parasites, as pathogen identification was based solely on previously known species or genera that are likely to cause human or animal diseases. More distantly related bacteria and fungi were not classified as pathogens, such that their actual diversity is greater than reported here. Additionally, our focus on only a subset of organ types restricts the full evaluation of zoonotic potential. For example, more eukaryotic pathogens are likely to be identified in the liver, including the Chinese liver fluke that is endemic in Guangdong province ^64–66^. Lastly, the geographic scope of the study was restricted to Guangdong province, and future research should extend to other regions of China for a broader understanding. As we expand the geographic regions, sampling size, and organ types, we expect to better assess both the zoonotic potential and the ecological correlates of the pathogens carried by these important mammalian disease reservoirs.

## Methods

### Sample collection

Samples from small mammals, comprising rodents (Rodentia) and shrews (Eulipotyphla), were collected in Guangdong province, China, between 2021 and 2022. Sampling was conducted across nine regions—Zhanjiang, Anpu, Maoming, Foshan, Heyuan, Jieyang, Shaoguan, Shenzhen, and Yunfu—each representing different geographical areas and natural habitats within the province. In most regions, collection occurred primarily during the late autumn and winter. However, in Anpu and Zhanjiang, sampling was conducted over the entire 12-month period of 2022. Animals were captured using baited cages in both agricultural and residential areas, then euthanized and dissected. Tissue samples, including lung, spleen, and gut, were harvested, immediately preserved in RNA Stabilization Solution (ThermoFisher, USA), and stored on dry ice before being transferred to a −80°C freezer.

All protocols for sample collection and processing were reviewed and approved by the Ethics Committee of Sun Yat-sen University (SYSU-IACUC-MED-2021-B0123)

### Sample processing, RNA extraction and sequencing

RNA extraction and sequencing were performed on 2,408 tissue samples from 858 individual animals. Each sample was homogenized in 600 μl of lysis buffer using a TissueRuptor (Qiagen, Germany), followed by total RNA extraction with the RNeasy Plus Mini Kit (Qiagen, Germany) according to the manufacturer’s protocol. Sequencing libraries were prepared using the MGIEasy RNA Library Prep Kit V3.0 (BGI, China). In brief, RNA was fragmented, reverse-transcribed, and converted into double-stranded cDNA. Unique dual-indexed cDNA molecules were circularized, and rolling-circle replication was employed to generate DNA nanoball (DNB)-based libraries. These libraries were then sequenced on the DNBSEQ T series platform (MGI, China), producing 150-bp paired-end metatranscriptomic reads. The target yield for each sample was 50 Gbp.

### Processing of sequencing data

For each sequence data set, the majority of ribosomal RNA (rRNA) reads were initially removed using URMAP (version 1.0.1480) ^67^. Adapters, duplicate and low-quality reads were filtered out using fastp (version 0.20.1, parameters: -q 20, -n 5,-l 50,-y, -c, -D) ^68^. The reads with low complexity were removed using PRINSEQ++ (version 1.2, options: -lc_entropy=0.5 -lc_dust=0.5) ^69^. Residual rRNA reads were further eliminated by mapping to the SILVA rRNA database (Release 138.1) ^70^ using Bowtie2 ^71^. Unless otherwise specified, all software was run with default settings.

### Molecular identification of host species

The identification of small mammal species was based on *de novo* assembled contigs containing COX1 gene sequences. For each sample, open reading frames (ORFs) from the assembled contigs were extracted using ORFfinder ^72^ and compared to COX1 reference sequences from the NCBI RefSeq database using the blastn program (version 2.14.1) with an e-value threshold of 10^-10^. To ensure accuracy, the core COX1 domain (cd01660) was confirmed using rpsblast against the Conserved Domain Database (CDD). Reads were subsequently mapped back to the assembled COX1 sequences to remove assembly error. To finalize species assignments, a phylogenetic tree incorporating COX1 sequences from this study, along with representative related sequences, was estimated using PHYML 3.0 (nucleotide substitution model: GTR+I+F+Γ_4_, with SPR branch-swapping) ^73^.

### Discovery of viruses and viral pathogens

The remaining clean non-rRNA reads were assembled into contigs using MEGAHIT (version 1.2.8) ^74^ with default settings and a minimum contig length of 300 bp. Assembled contigs were then searched against the NCBI nr database using DIAMOND blastx (version 2.0.14) ^75^ with an e-value cutoff of 10^-5^ to balance high sensitivity and reduce false positives. Contigs were provisionally categorized based on the NCBI taxonomy of the best-matching protein, and viral-related contigs were extracted. Host-related regions in the viral contigs were removed by aligning them against the NCBI RefSeq genome database using the blastn program ^76^ with an e-value cutoff of 10^-10^. Viral identities were further confirmed by checking for the presence of specific marker genes: the RNA-dependent RNA polymerase (RdRP) for RNA viruses, non-structural protein 1 (NS1) for *Parvoviridae*, hexon for *Adenoviridae*, capsid protein (CP) for *Circoviridae*, and ORF1 protein for *Anelloviridae*. These marker proteins were then aligned and examined manually to ensure they contained the conserved motif(s) of the corresponding protein. New virus species were determined according to the species demarcation criteria established by the International Committee on Taxonomy of Viruses (ICTV). To identify viruses associated with mammalian hosts, phylogenetic analyses were conducted for each virus supergroup (i.e., phylum or class level taxonomic groups). Only viral contigs that clustered within families known to infect mammals were classified as mammalian viral pathogens and retained for further analysis.

### Discovery of bacterial and eukaryotic pathogens

The remaining contigs were screened against the Conserved Domain Database (CDD) using the rpsblast program ^76^, with an e-value cutoff of 0.01. We targeted contigs containing specific marker genes for identifying eukaryotic microbes, specifically COX1 (cd01663) and EF1a (cd01883), and bacteria, specifically ftsY (TIGR00064), GroEL (TIGR02348), nusG (TIGR00922), rplA (TIGR01169), rplC (TIGR03625), and rpoB (TIGR02013). To facilitate taxonomic identification and to remove false positives, the sequences of these marker genes were then compared against the nt and nr databases with e-values set to 10^-10^ and 10^-5^, respectively. For bacterial contigs, reads were mapped back to the homologous gene from the closest relative when applicable, or to the relevant contig if more distantly related, using Bowtie2 in ‘end-to-end’ mode ^71^. Species identification for both bacterial and eukaryotic microbes was then conducted through phylogenetic analyses involving these marker genes. Pathogenic microbes were identified based on their relationship to known bacterial and eukaryotic pathogens at the species and genus level.

### Quantification of pathogen genomes/transcriptomes

To estimate pathogen abundance, reads were mapped to pathogen genomes (viruses and bacteria) or to a set of marker genes (for eukaryotes) using Bowtie2 (with end-to-end alignment). Pathogen abundance was measured as the number of reads mapped per million non-rRNA reads (RPM). Two criteria were applied to reduce potential false positives. First, index-hopping, which can occur during high-throughput sequencing when reads are misassigned between samples, was filtered out using the following rule: if the total read count for a specific virus in a given library was less than 0.1% of the highest read count for that virus in the same sequencing lane, it was considered a false positive due to index-hopping. Second, low-abundance pathogens (RPM < 1) and those with low genome or gene coverage (i.e., less than 300 base pairs) were also likely to be false positives and were excluded ^37,77^.

### Phylogenetic analyses

To determine the taxonomy of newly identified pathogens, representative marker proteins or genes related to those identified in this study were downloaded from NCBI/GenBank. Phylogenetic trees were estimated at the genus or family level. Sequences were first aligned using the L-INS-i algorithm in MAFFT (version 7) ^78^, and ambiguously aligned regions were removed using TrimAl ^79^. Maximum likelihood (ML) trees were then inferred using PhyML (version 3.0) ^73^, with the GTR substitution model used for nucleotide sequence alignments and the LG model used for amino acid sequence alignments, with the SPR branch-swapping algorithm employed to find the optimal tree topology.

### Collection and processing of environmental data for epidemiological analyses

To assess how environmental factors shape pathogen diversity and composition, we collected climate, mammal richness, and land-use data for each sampling location from publicly available sources. Climate data were obtained from TerraClimate ^80^, utilizing 14 variables to evaluate their influence on rodent pathogens. Definitions of these variables are available on the TerraClimate website (https://www.climatologylab.org/terraclimate.html). To address co-linearity among the climate variables, we performed principal component (PC) analysis. The first three PCs—CPC1, CPC2, and CPC3—were used in subsequent statistical analyses, explaining 57.96%, 16.15%, and 11.40% of total variance, respectively (cumulatively 85.51%). Based on the projection lengths of raw bioclimatic variables onto these PCs, we interpreted the components as follows:

i. CPC1 primarily reflects negative correlations with temperature, shortwave radiation, evapotranspiration, and precipitation. Lower CPC1 values indicate higher temperatures and greater water evaporation.
ii. CPC2 is mainly associated with wind and evapotranspiration. Higher CPC2 values suggest increased water evaporation and lower wind speeds, indicating a harsher thermal environment.
iii. CPC3 captures precipitation variability, with higher values indicating greater fluctuation in precipitation levels.

Land-use data were sourced from the CNLUCC database ^81^, while NDVI values were derived from publicly available remote sensing satellite data. Mammal richness data were obtained from the IUCN database ^82^.

### Statistical methods

All statistical analyses were conducted using R version 4.3.1.

### Assessing environmental and host factors influencing pathogen species richness

To investigate how environmental and host factors influence pathogen species richness, we applied generalized linear models using a negative binomial regression. The factors considered in the analysis included rodent species identity, environmental characteristics, date, and region of sample collection. Environmental characteristics comprised three principal components as described above (CPC1, CPC2, and CPC3), NDVI, and mammal richness. Model selection was performed based on Akaike Information Criterion (AIC), evaluating all possible combinations of variables using the MuMIn package in R. The contribution of each variable to model performance was determined by comparing the deviance explained by the full model to that of models where individual variables were removed.

### Analysis of pathogen composition and cross-species transmission

We quantified the pathogens shared among animal species and visualized the results using the ComplexUpset package ^83^. The pathogen-sharing network was initially constructed using the ggraph package ^84^ in R, refined through manual adjustments, and visualized using the ggplot2 package. In addition, we investigated the factors influencing pathogen composition (or number of shared pathogens between individual animals) by applying generalized linear models (GLMs), considering host phylogenetic distance, climate variations (Euclidean distance), land use variations, and spatial distance. The effect of each factor was quantified through a model selection process similar to that used in earlier analyses.

## Data availability

All sequencing data have been submitted to the China National GeneBank (CNGB) Sequence Archive with BioProject ID: CNP0004378. The genomic sequences of pathogens generated in this study will be deposited in the NCBI GenBank database or China National GeneBank DataBase and is temprorily available at Figshare for reviewing purposes: https://doi.org/10.6084/m9.figshare.28054424.

## Acknowledgements

This work was supported by the National Key Research and Development Program of China (2024YFC2607502 and 2021YFC2300905), Natural Science Foundation of Guangdong Province of China (2022A1515011854), National Natural Science Foundation of China (82341118 and 32270160), Shenzhen Science and Technology Program (KQTD20200820145822023), Major Project of Guangzhou National Laboratory (GZNL2023A01001), Open Project of BGI-Shenzhen Shenzhen 518000, China (BGIRSZ20210001), Guangdong Province “Pearl River Talent Plan” Innovation, Entrepreneurship Team Project (2019ZT08Y464), Guangdong Provincial Key Laboratory of Pathogen Detection for Emerging Infectious Disease Response (2023B1212010010) and the Fund of Shenzhen Key Laboratory (ZDSYS20220606100803007). E.C.H. was supported by an NHMRC (Australia) Investigator Award (GNT2017197) and by AIR@InnoHK administered by the Innovation and Technology Commission, Hong Kong Special Administrative Region, China. We gratefully acknowledge colleagues at BGI-Shenzhen and the China National Genebank (CNGB) for library construction, and sequencing.

## Author contributions

Conceptualization, X.Z., E.C.H., and M.S.; Methodology, G.-Y.X., D.-X.W., X.Z., W.-C.W. and M.S.; Investigation, G.-Y.X., D.-X.W., X.Z., Q.-Q.C., H.-L.Z., W.-Q.Z, P.-B.S, Q.-Y.G., J.-B.K., M.-W.P., W.-C.W. and M.S.; Sample processing, G.-Y.X., M.-W.P., Y.-Q.L., J.W, S.-J.L., J.-X.C., G.-Y.L.,X.H., Q.-Y.G., J.-B.K., H.-L.Z., P.-B.S, Z.-R.R., W.-Q. Z., J.-H.L., P.-H.J. and G.-P.K.; Writing – Original Draft, G.-Y.X. and M.S.; Writing – Review and Editing, All authors; Funding Acquisition, D.-X.W., X.Z., Z.-Q.D., and M.S.; Resources (sampling), G.-Y.X., M.-W.P., Y.-Q.L., Q.-Q.C., A.C., Z.P., D.-Q.L., S.-P.G., J.-J.L., W.-Q.L., J.W., J.-J.L., C.-Y.S. and J.-D.C.; Resources (Computational), D.-X.W., J.-H.L., Z.-Q.D., W.-C.W. and M.S.; Supervision, D.-X.W., X.Z., Z.-Q.D. and M.S..

## Supplementary Figures

**Supplementary Figure 1. RNA and DNA viruses in small mammals.** Maximum likelihood phylogenetic trees were estimated at the supergroup (RNA viruses) or family (DNA viruses) level based on conserved viral proteins, namely RNA-dependent RNA polymerase (RdRP) for RNA viruses, non-structural protein 1 (NS1) for *Parvoviridae*, hexon protein for *Adenoviridae*, capsid protein for *Circoviridae*, and ORF1 protein for *Anelloviridae*. All trees were midpoint-rooted for clarity. Vertebrate-associated virus clusters are highlighted in blue. Solid circles indicating viruses identified in this study, which are colored according to vertebrate association.

**Supplementary Figure 2. Identification of bacterial pathogens.** Each maximum likelihood phylogenetic tree represents the diversity of a bacterial genus and was estimated using the groEL chaperonin (GroEL) gene. All trees were midpoint-rooted for clarity. The species names of each sequence are shown to the right of the tree. Solid circles on the tree indicate bacteria identified in this study, colored according to their potential pathogenicity in mammals.

**Supplementary Figure 3. Identification of eukaryotic pathogens.** Maximum likelihood phylogenetic trees were estimated using the elongation factor 1-alpha (EF1a) or COX1 genes. All trees were midpoint-rooted for clarity. Solid circles on the trees indicate pathogens identified in this study.

**Supplementary Figure 4. Phylogeny of 11 vertebrate-associated RNA viral families identified in this study**. All trees were midpoint-rooted for clarity. The taxonomy for each tree is shown in the top-left corner. Red solid circles on the trees represent RNA viruses identified in this study.

**Supplementary Figure 5. Phylogeny of three vertebrate-associated DNA viral families identified in this study.** All trees were midpoint-rooted for clarity. The taxonomy for each tree is shown in the top-left corner. Red solid circles on the trees represent RNA viruses identified in this study.

**Supplementary Figure 6. Distribution and abundance levels of 10 cross-order transmission pathogens in the sampled individuals.** The relative abundance of viruses in each library was calculated by RPM. Hosts are represented by different colors.

## Supplementary tables

Supplementary Table 1. Sample information.

Supplementary Table 2. Library information.

Supplementary Table 3. Pathogen information.

